# Molecular motors orchestrate pause-and-run dynamics to facilitate intracellular transport

**DOI:** 10.1101/2025.11.22.689936

**Authors:** Yusheng Shen, Kassandra M. Ori-McKenney

**Affiliations:** Department of Molecular and Cellular Biology, University of California, Davis, Davis, CA 95616, USA

**Keywords:** Intracellular Transport, Molecular Motors, Kinesin, Dynein, Vesicle Trafficking, Microtubules, Motor Engagement, Cytoplasmic Diffusion, Single-Particle Tracking

## Abstract

Intracellular transport is essential for cellular organization and function. This process is driven by molecular motors that ferry cargo along microtubules, but is characterized by intermittent motility, where cargoes switch between directed runs and prolonged pauses. The fundamental nature of these pauses has remained a mystery, specifically whether they are periods of motor detachment and passive drifting or states of active motor engagement. By combining single-particle tracking with largescale motion analysis, we discovered that pauses are not passive. Instead, they are active, motor-driven states. We uncovered a unifying quantitative law: the diffusivity of a vesicle during a pause scales with the square of its velocity during a run. This parabolic relationship, *D*_eff_ ∝ *v*^2^, holds true for both kinesin and dynein motors, different cargo types, and a variety of cellular perturbations. We show that this coupling arises because the number of engaged motors governs motility in both states. When we reduce motor engagement, either by severing the motor-cargo link or by chemically modifying the microtubule track, vesicles move slower and become trapped in longer, less mobile pauses, failing to reach their destination. Our work redefines transport pauses as an essential, motor-driven part of the delivery process, revealing a fundamental principle that ensures robust cargo transport through the crowded cellular environment.

## Introduction

Mammalian cells expend considerable energy to transport materials through their crowded cytoplasm. This active transport is powered by molecular motors, such as kinesins and cytoplasmic dynein, which move cargoes directionally along microtubule tracks to maintain cellular organization and homeostasis (1). By converting chemical energy into mechanical work, these motors enable long-range transport that overcomes the limits of passive diffusion, particularly for large cargoes such as vesicular organelles and ribonucleoprotein complexes in the dense, viscoelastic cytoplasm (2– 5). To operate efficiently, motor–cargo complexes must adapt to a dynamic and complex intracellular landscape shaped by physical obstacles (6), membrane-membrane contacts (7, 8), microtubule-associated proteins (MAPs) (9–13), and tubulin post-translational modifications (PTMs) (14–18), all of which modulate motor activity.

Perhaps due to this complexity, a hallmark of intracellular transport is intermittency. Cargoes including endosomes, lysosomes, secretory vesicles, mitochondria, and autophagosomes switch between short, directed runs and longer, seemingly stalled, diffusive pauses (19–28). These pauses can consume up to 80% of a cargo’s travel time, presenting a paradox. While runs clearly reflect active motor-driven motion, the fundamental nature of pauses remains debated. This contrasts sharply with the behavior of individual cargo-free motors in vitro, which move processively along bare microtubules (11, 29, 30). Why, then, do cargoes in cells spend most of their time in a seemingly inefficient paused state?

In cells, pauses have been attributed to various causes, including physical barriers (6, 23, 31), transient membrane tethers (8, 22, 32), regulation by MAPs and tubulin PTMs, and mechanical “tug-of-war” between opposing motors (12, 20, 21, 26, 33, 34). However, a critical and unresolved question remains: during a pause, are motor-cargo complexes actively engaged with the microtubule, withstanding viscoelastic cytoplasmic forces, or do they detach from the microtubule and passively diffuse? Directly distinguishing these possibilities in living cells has been challenging because it requires resolving the tripartite interactions of microtubules, motors, and cargoes with both high spatial and temporal resolution coupled with large-scale statistical power to analyze highly heterogeneous behaviors (23, 28). Resolving this is essential for understanding how cells precisely allocate energy to enable long-range intracellular transport.

To address this question, we combined single-particle tracking with large-scale statistical motion-state analysis of cargo-free kinesin-1 and its physiological cargo, Rab6A-positive secretory vesicles. We altered cytoplasmic fluidity, motor–cargo coupling, and motor–microtubule interactions and found that vesicle mobility during pauses is governed by motor activity in the run state and is independent of cytoplasmic viscosity. Parallel analysis of dynein-driven retrograde transport of Rab5-positive early endosomes revealed the same coordinated relationship, indicating a conserved mechanism between transport directions. Together, our live-cell observations uncover a robust quantitative coupling between pause-state diffusivity and run-state velocity, *D*_eff_ ∝ *v*^2^, that holds across motor families, perturbations, and cargo types. This unifying principle demonstrates that motor engagement directly coordinates both motility states, thereby ensuring robust cargo delivery through the complex intracellular landscape.

## Results

### Vesicle-bound motor complexes exhibit distinct transport dynamics compared to cargo-free kinesin-1 motors

To investigate the origins of pausing during intracellular transport, we compared the movement of cargo-free kinesin1 motors to their physiological cargo, secretory vesicles labeled with EGFP-Rab6A. These vesicles are primarily driven anterogradely by kinesin-1 and kinesin-3 motors (24, 25). While kinesin-1 moves processively along bare microtubules in vitro (Fig.1A), vesicle cargoes pause frequently in cells, raising the question about the nature of these pauses (10, 30, 35). To perform live single-molecule imaging of kinesin-1 in BEAS-2B cells, we tagged the truncated kinesin-1 motor, KHC(1-560), with the bright and stable fluorescent protein, mStaygold (36), and transiently expressed it at low levels using a truncated cytomegalovirus (CMV) promoter (Fig. 1B) (37). This construct lacks the cargo-binding domain of kinesin-1, allowing us to directly examine how the in vivo microtubule landscape influences the motility of the active dimeric motor KHC(1–560) (1, 38). In BEAS-2B cells, single KHC(1–560)–mStayGold motors exhibited unidirectional, processive motility along linear tracks without pausing, showing some preferential movement along specific microtubules, consistent with previous findings (39, 40). By analyzing over 2000 motor tracks, we found that KHC(1–560) exhibited a single characteristic run time of *∼*0.9 s on in vivo microtubules, despite the heterogeneous landscape of MAPs and tubulin PTMs (25). This suggests that single cargo-free kinesin-1 motors move rather freely along microtubules in cells, similar to their behavior on bare microtubules in vitro.

**Fig. 1.**
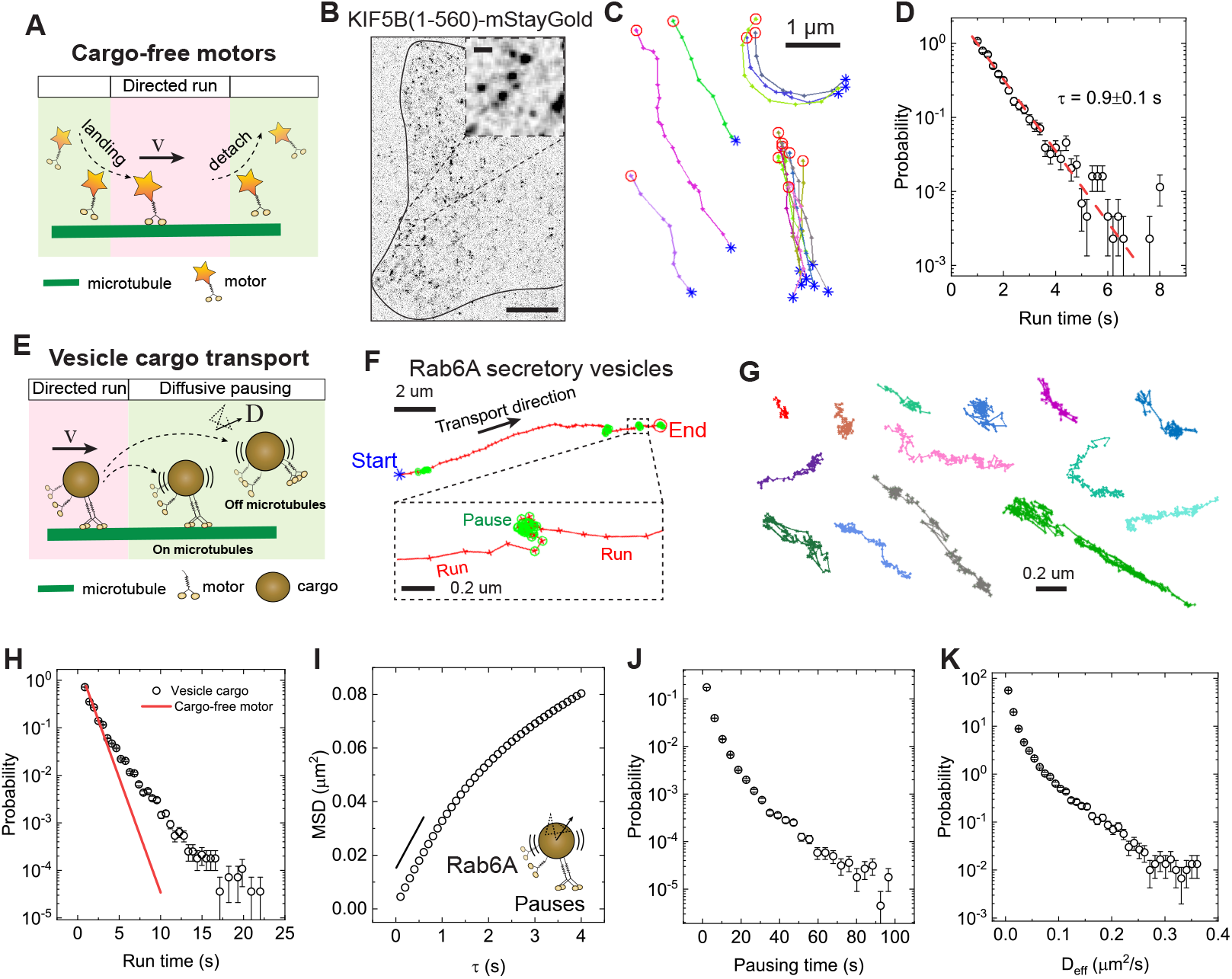
Pausing state of intracellular vesicle transport is diffusive-like and vastly heterogeneous. (A) Schematic illustration of the landing, directed run and detach dynamics of a cargo-free KIF5B motor. (B) Image of a BEAS-2B cell expressing KIF5B(a.a. 1-560)-mStayGold visualized by TIRF microscopy. Scale bar: 10 *µ*m. Inset, magnified view of KIF5B motors. Scale bar: 1 *µ*m. (C) Representative trajectories (colored lines) of truncated KIF5B motors (5 fps for 3 min). Blue stars and red circles mark the start and end point of the trajectory, respectively. Scale bar: 1 *µ*m. (D) Measured probability density function (PDF) of trajectory-based run time, *T*_*p*_, for truncated KIF5B motors expressed in BEAS-2B cells. The measured *T*_*p*_ could be fitted to a simple exponential distribution (red dashed line), with a characteristic *T*_*p*_ = 0.9 *±* 0.1s. Total trajectories analyzed from the mobile fraction are n = 2183 from 16 cells. (E) Schematic illustration of the intermittent switching between directed run and diffusive pausing of secretory vesicles driven by teams of motors. (F) Magnified trajectory image showing a Rab6A-positive vesicle from BEAS-2B cells. Blue stars and red circles mark the start and end point of the trajectory, respectively. Pause states are highlighted by green circles. Motion states were detected and extracted by a homemade algorithm. Scale bar: 2 *µ*m. Inset shows a magnified view of the pause state. Scale bar: 200 nm. (G) Representative pausing segments extracted from mobile trajectories. Scale bar: 200 nm. (H) Measured PDF of the dwell time for run segments extracted from mobile trajectories (black circles). Run time of single KIF5B-motors are replotted for comparison (red line). (I) Measured mean squared displacement (MSD) as a function of delay time *τ* for all the pausing segments extracted from the mobile trajectories of Rab6A-positive vesicles. (J) Measured PDF of pausing time for Rab6A-positive vesicles. (K) Measured PDF of the effective diffusion coefficient *D*_eff_ in the pausing state for Rab6A-positive vesicles. Total trajectories analyzed from the mobile fraction are n = 30418 from n = 48 cells.

Given that single motors do not pause, we next asked why vesicles driven by multiple motors pause so frequently. We used a previously established algorithm to extract distinct motion states from vesicle trajectories (25). For Rab6Apositive secretory vesicles in BEAS-2B cells, pauses occurred randomly along microtubules and accounted for up to 60% of the total travel time (Fig. 1E-G). Analysis of over 51,000 run segments from Rab6A-vesicle trajectories revealed that their run-time distribution was not a simple exponential and was significantly longer than that of single cargo-free KHC(1–560) motors (Fig. 1H). This is consistent with transport being driven by multiple motors (24, 41). The pausing states themselves were characterized by confined, jiggling movements and some pauses exhibited a distinct, strip-like pattern aligned with microtubules (Fig. 1G). These pauses exhibited diffusive-like behavior at short timescales (*<*1 s), with mean square displacements (MSDs) increasing linearly with delay time (*τ* ), before transitioning to confined motion at longer timescales (Fig. 1I). The pausing behavior was highly heterogeneous, with both pausing duration and diffusivity varying more than 50-fold between events (Fig. 1J-K), suggesting that diverse cellular mechanisms likely regulate these pauses.

### Rab6A vesicle pauses are microtubule-bound states coupled to motor activity

Since the pauses during vesicle transport displayed a strong microtubule-associated pattern (Fig. 1G), we next sought to determine whether these pauses represented states in which the motor–vesicle complex remained associated with, or became detached from, microtubules (Figs. 1E and 2A). If pauses represent detached states, vesicle mobility should positively correlate with cytoplasmic fluidity, similar to the behavior of inert particles diffusing in the cytoplasm (Fig. 2A) (42). To test this hypothesis, we exposed cells to extracellular osmotic environments ranging from hypoosmotic to hyperosmotic conditions (250–400 mOsm) in order to either increase or decrease cytoplasmic fluidity (25). We then quantified these fluidity changes by tracking the motion of 40-nm-diameter genetically encoded multimeric (GEM) nanoparticles in BEAS-2B cells (Figs. 2B-D). Consistent with previous findings (25, 43), GEM diffusivity increased under hypotonic treatment and decreased under hypertonic treatment (Figs. 2B-D), confirming that cytoplasmic fluidity was modulated accordingly.

**Fig. 2.**
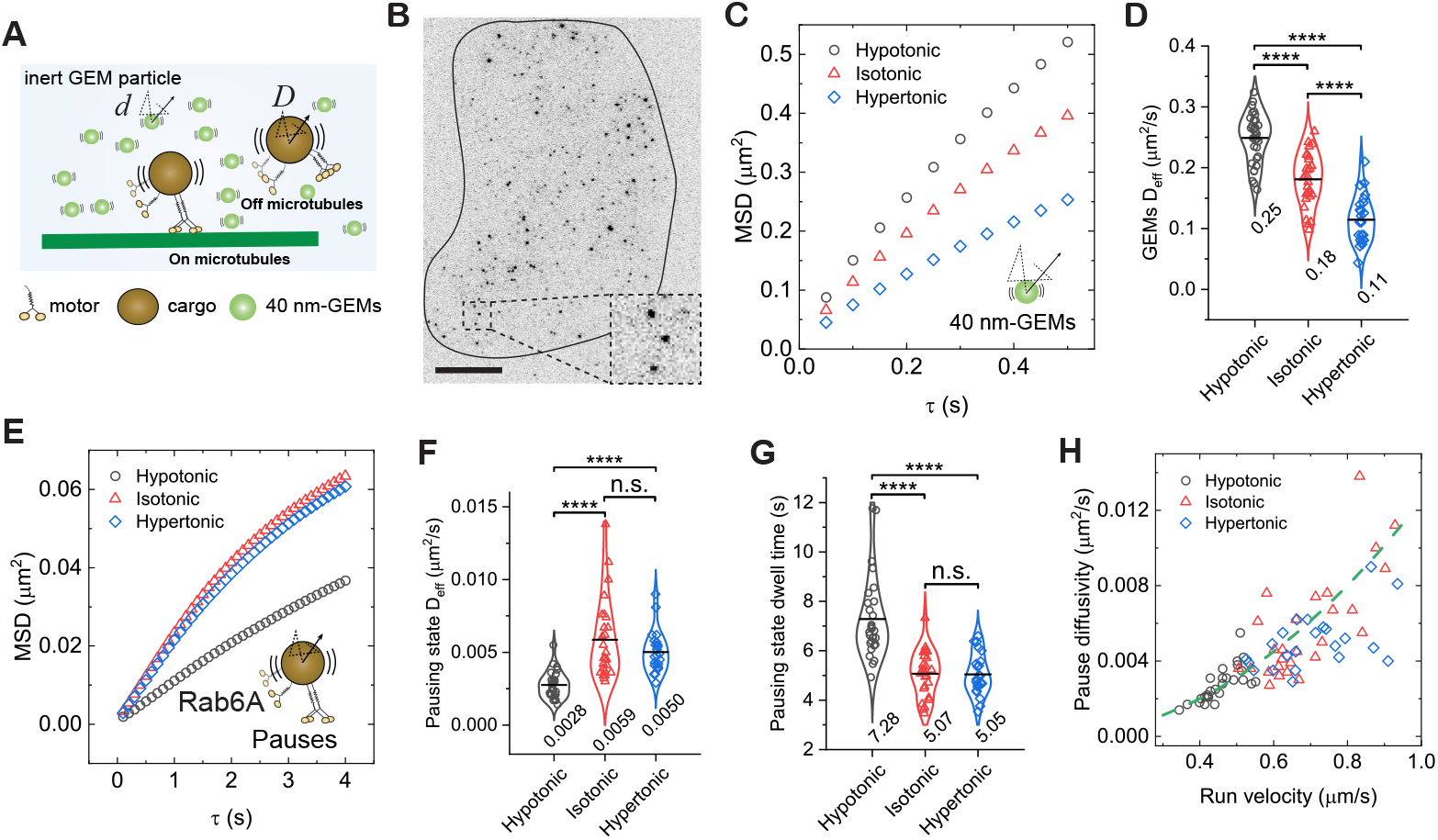
Diffusivity of Rab6A-positive vesicles in the pausing state positively correlates with motor activity in the run state but not cytoplasmic fluidity. (A) Schematic illustration of the diffusion of inert 40 nm-GEMs and diffusive-like motion of Rab6A-positive vesicles in the pausing state either on microtubules or off microtubules. (B) Image of a BEAS-2B cell expressing 40-nm GEMs visualized by TIRF microscopy. Inset: magnified view of GEMs. Scale bar, 10 *µ*m. (C) Measured MSDs of GEMs as a function of delay time *τ* for cells treated with isotonic (310 mOsm), hypotonic (250 mOsm), or hypertonic (400 mOsm) solutions. (D) Measured *D*_eff_ of GEMs for cells treated with different osmotic solutions. \*\*\*\**p <* 0.0001. Total trajectories analyzed from the mobile fraction are n = 361318, 196137 and 96875 from n = 33, 29 and 27 cells from 3 independent experiments. (E) Measured MSDs of Rab6A-positive vesicles in the pausing state as a function of delay time *τ* for cells treated with different osmotic solutions. (F and G) Measured *D*_eff_ of Rab6A-positive vesicles in the pausing state (F) and pausing time (G) for cells treated with different osmotic solutions. \*\*\*\**p <* 0.0001, n.s. indicates not significant. Total trajectories analyzed from the mobile fraction are n = 29688, 32248 and 22572 from n = 42, 40 and 34 cells from 3 independent experiments. (H) Measured pausing state diffusivity as a function of run-state velocity for Rab6A-positive vesicles from each cell. The green dashed line represents a fit to a parabolic distribution curve, given by, *D*_eff_ = 0.0125*v*^2^. All violin graphs display all data points with means. P-values were calculated using a student’s t-test.

We next investigated the effect of cytoplasmic fluidity on the pausing mobility of Rab6A-positive vesicles. Unexpectedly, hypotonic treatment, which increased cytoplasmic fluidity, reduced vesicle diffusivity during pauses, whereas hypertonic treatment, which decreased cytoplasmic fluidity, had no detectable effect compared with the isotonic control (Figs. 2E–F). Notably, the reduced diffusivity under hypotonic conditions correlated with longer vesicle dwell times in the pausing state (Fig. 2G). The response of pausing behavior to changes in cytoplasmic fluidity correlated with that of run behavior reported previously for Rab6A-positive vesicles, where hypotonic treatment inhibited directed runs (25). To further define the relationship between the two states, we plotted pause-state diffusivity against run-state velocity for Rab6A-positive vesicles. Strikingly, all data from different fluidity conditions collapsed onto a single parabolic curve (Fig. 2H), revealing a robust positive correlation between pausing mobility and run velocity that is independent of cytoplasmic fluidity. These results indicate that the motor–vesicle complex remains associated with microtubules during pauses, and that reduced motility in the run state is coupled with reduced mobility in the pausing state, leading to prolonged trapping of vesicles in the pausing state.

### Kinesin-1 engagement governs vesicle motility in run and pause states

To directly examine the role of kinesin-1 in coordinating vesicle motility during run and pause states, we treated cells with kinesore, a small-molecule compound that has a dual effect of activating kinesin-1 on microtubules while simultaneously disrupting its association with native cargoes (44, 45), like Rab6A-positive vesicles (Fig. 3A). While this severs the specific kinesin-1-cargo linkage, the net effect on the vesicle is a reduction in the number of processive, cargo-bound motors capable of generating cooperative force. Treating BEAS-2B cells with 100 *µ*M kinesore for 1 hour did not noticeably alter the radial microtubule network (Fig. 3B). As predicted, acute kinesore treatment impaired the run motility of Rab6Apositive vesicles, significantly reducing all measured run parameters compared with control (Figs. 3C-G). This confirms that kinesore disrupts the engagement of kinesin-1 motors with the cargo.

**Fig. 3.**
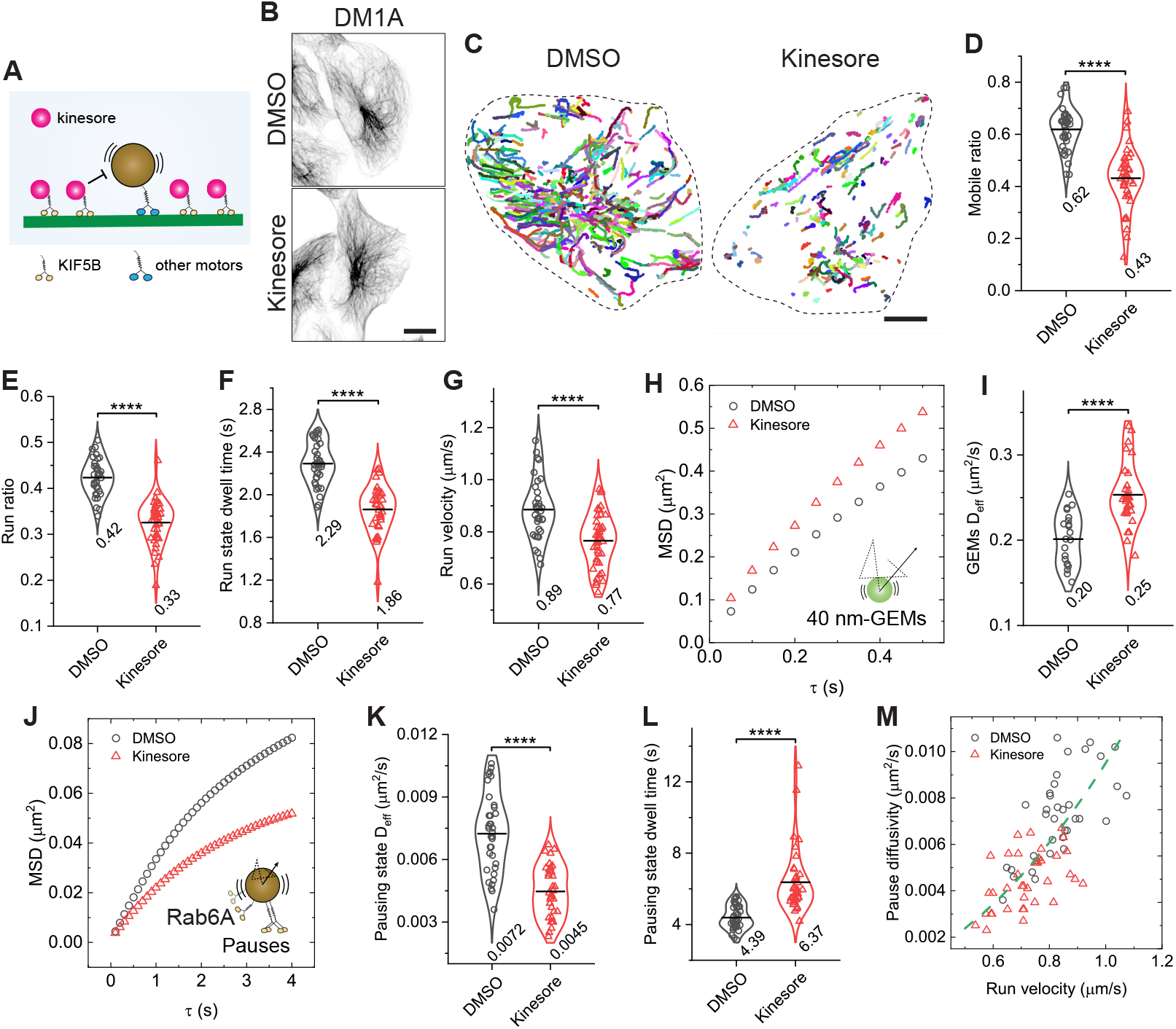
Reducing active motors on vesicles through kinesore treatment decreases pausing state diffusivity. (A) Schematic illustration showing the effect of Kinesore on kinesin-1 activity: Kinesore promotes kinesin-1 binding to microtubules while inhibiting its interaction with native vesicle cargoes. (B) Representative images comparing the microtubule network in BEAS-2B cells treated with 100 *µ*M Kinesore or vehicle control (DMSO). Scale bar: 20 *µ*m. (C) Representative trajectories of the mobile fraction of Rab6A-positive vesicles for cells treated with DMSO or 100 *µ*M Kinesore. Scale bars, 10 *µ*m. (D-G) Measured run parameters of Rab6A-positive vesicles in BEAS-2B cells treated with DMSO or Kinesore. \*\*\*\**p <*0.0001. (H) Measured MSDs of GEMs as a function of delay time *τ* for cells treated with DMSO or 100 *µ*M Kinesore. (I) Measured *D*_eff_ of GEMs for cells treated with DMSO or Kinesore. \*\*\*\**p <*0.0001. Total trajectories analyzed from the mobile fraction are n = 74770 and 39858 from n = 21 and 28 cells from 2 independent experiments. (J) Measured MSDs of Rab6A-positive vesicles in the pausing state as a function of delay time *τ* for cells treated with DMSO or Kinesore. (K and L) Measured *D*_eff_ of Rab6A-positive vesicles in the pausing state (K) and pausing time (L) for cells treated with DMSO or Kinesore. \*\*\*\**p <*0.0001. Total trajectories analyzed from the mobile fraction are n = 29467 and 16413 from n = 36 and 40 cells from 3 independent experiments. (M) Measured pausing state diffusivity as a function of run-state velocity for Rab6A-positive vesicles from each cell. The green dashed line represents a fit to a parabolic distribution curve, given by, *D*_eff_ = 0.0095*v*^2^. All violin graphs display all data points with means. P-values were calculated using a student’s t-test.

We next asked if reducing active kinesin-1 motors on vesicles similarly affects the pausing state. Despite kinesore increasing cytoplasmic fluidity (Figs. 3H–I), Rab6A-positive vesicle diffusivity during pauses was reduced, and trapping times were increased (Figs. 3J-L). Kinesore-induced increases in cytoplasmic fluidity may result from enhanced microtubule remodeling, which could actively agitate the surrounding cytoplasm (44, 46). Nonetheless, the correlation between pause-state diffusivity and run-state velocity mirrored that observed in cells under varying osmotic conditions (Fig. 3M), suggesting that mobility in both states is intrinsically linked to active kinesin-1 simultaneously engaging with vesicle cargo and microtubules.

### Microtubule detyrosination coordinates vesicle motility in run and pause states

Having established that disrupting motor–cargo interactions with kinesore impairs both run and pause states, we next asked if a similar effect would occur by perturbing the motor’s engagement with the microtubule track itself. We focused on microtubule detyrosination, a posttranslational modification that has been shown to efficiently inhibit the landing and processivity of the kinesin-3 motor, KIF13B, in vitro by interfering with its CAP-Gly domain (18). Given KIF13B’s role in transporting Rab6A-positive secretory vesicles (24), we tested if detyrosination would disrupt the coordination of pause-run motility at the motor-microtubule interface (Fig. 4A).

**Fig. 4.**
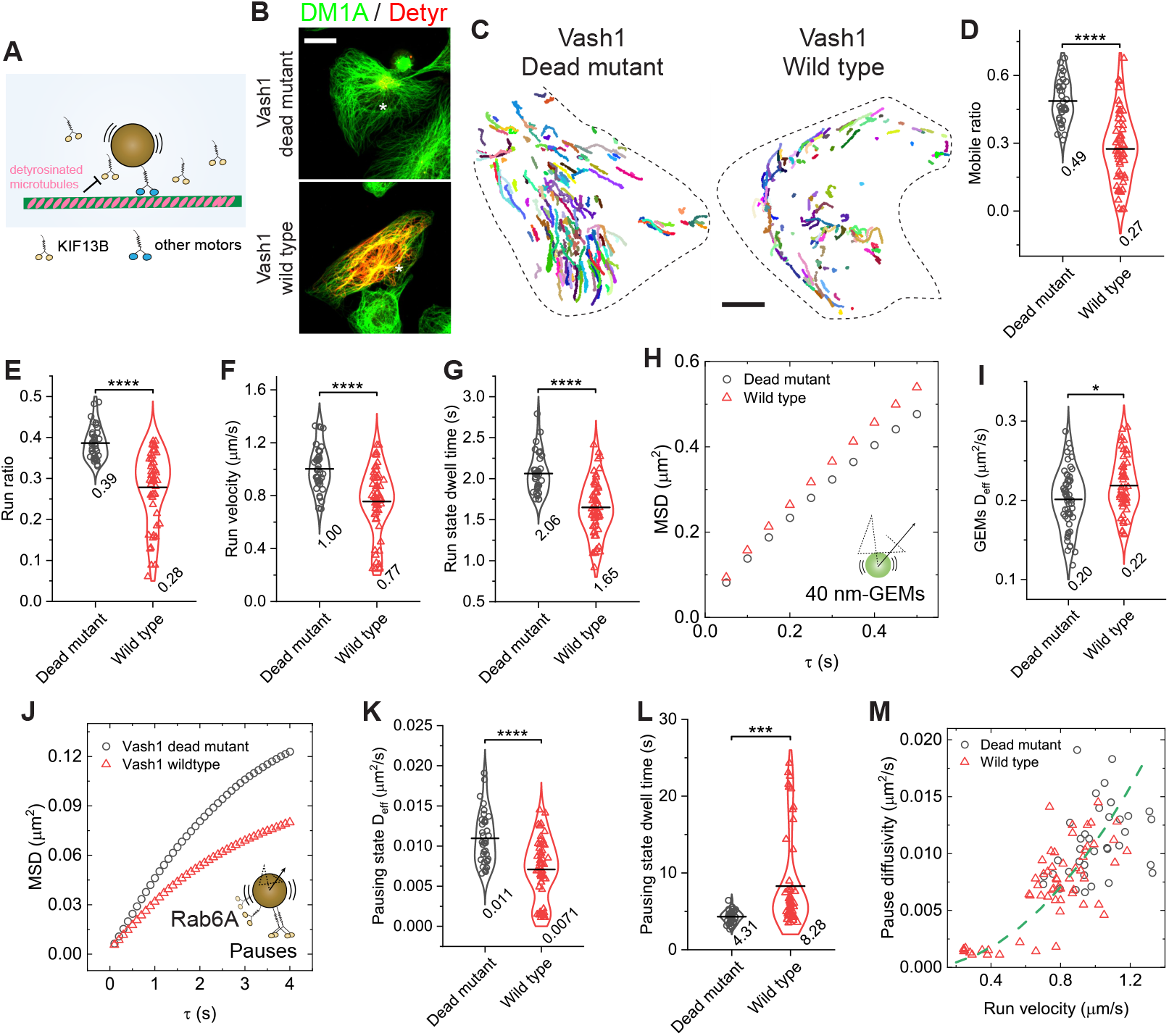
Microtubule detyrosination inhibits anterograde transport in both run and pausing states. (A) Schematic illustration of inhibition of kinesin-3 activity by microtubule detyrosination. (B) Representative images showing *α*-tubulin (green) and detyrosinated *α*-tubulin (red, Detyr) for BEAS-2B cells transiently expressing the wild type (wild type) or catalytically dead mutant (C168A, dead mutant) of VASH1-EGFP-P2A-SVBP. Scale bar, 20 *µ*m. (C) Representative trajectories of the mobile fraction of Rab6Apositive vesicles for cells transiently expressing the wild type or the dead mutant of VASH1. Scale bars, 10 *µ*m. (D-G) Measured run parameters of Rab6A-positive vesicles in BEAS-2B cells transiently expressing the wild type or the dead mutant of VASH1. \*\*\*\**p <*0.0001. (H) Measured MSDs of GEMs as a function of delay time *τ* for cells transiently expressing the wild type or the dead mutant of VASH1. (I) Measured *D*_eff_ of GEMs for cells transiently expressing the wild type or the dead mutant of VASH1. \**p*=0.019. Total trajectories analyzed from the mobile fraction are n = 138981 and 139802 from n = 57 and 63 cells from 4 independent experiments. (J) Measured MSDs of Rab6A-positive vesicles in the pausing state as a function of delay time *τ* for cells transiently expressing the wild type or the dead mutant of VASH1. (K and L) Measured *D*_eff_ of Rab6A-positive vesicles in the pausing state (K) and pausing time (L) for cells transiently expressing the wild type or the dead mutant of VASH1. \*\*\*\**p <*0.0001, \*\*\**p*=0.00014. Total trajectories analyzed from the mobile fraction are n = 15924 and 13421 from n = 43 and 59 cells from more than 3 independent experiments. (M) Measured pausing state diffusivity as a function of run-state velocity for Rab6A-positive vesicles from each cell. The green dashed line represents a fit to a parabolic distribution curve, given by, *D*_eff_ = 0.011*v*^2^. All violin graphs display all data points with means. P-values were calculated using a student’s t-test.

Detyrosination of *α*-tubulin is catalyzed by an enzyme complex composed of a vasohibin (VASH1 or VASH2) and a small vasohibin-binding protein (SVBP) (47, 48). Transient expression of wild-type VASH1 in BEAS-2B cells markedly increased microtubule detyrosination compared with the catalytic-dead mutant (C168A), consistent with our previous findings (Fig. 4B) (26). We found that increasing microtubule detyrosination drastically impaired the run motility of Rab6A-positive vesicles, consistent with previous in vitro results (Figs. 4C–G) (18). We then assessed the pausing state. Analysis of 40-nm GEM diffusion showed that increased microtubule detyrosination slightly increased cytoplasmic fluidity, in agreement with previous measurements (Figs. 4H–I) (26). Strikingly, despite the more fluid environment, vesicles on detyrosinated microtubules showed substantially reduced diffusivity and prolonged trapping during pauses (Figs. 4J–L). This result mirrors the effect of kinesore treatment. The relationship between pause-state diffusivity and run-state velocity followed the same parabolic curve observed across perturbations (Fig. 4M). Together, these results demonstrate that impairing the motor-microtubule interaction coordinately regulates vesicle motility in both run and pause states. This finding, combined with our kinesore data, reveals that the effective number of motors productively engaged in transport is the key parameter governing this coupled behavior.

### Motor engagement coordinate pause–run dynamics of retrograde transport

Our findings support that the run-pause dynamics of kinesindriven vesicles are coordinated by motor engagement. We next asked whether a similar mechanism governs dyneindriven retrograde transport. To address this, we performed analogous analyses on Rab5-positive early endosomes, which are predominantly transported retrogradely by dynein–dynactin complexes in BEAS-2B cells (Fig.5) (22, 25, 49). Compared with kinesin-driven Rab6A-positive vesicles, Rab5-positive endosomes exhibited more frequent pauses, spending approximately 82% of their total transport time in the pausing state (Figs. 5A–B and S1). Similar to Rab6A vesicles, these pauses were diffusive-like and highly heterogeneous, with many trajectories displaying strip-like patterns indicative of microtubule association (Figs. S1 and 5B). We previously found that hypotonic treatment only modestly enhances Rab5-endosome run behavior, whereas hypertonic treatment downregulates it (25). Consistent with a motor-engaged pausing-state, the diffusivity of Rab5-positive endosomes during pauses did not scale with the increased cytoplasmic fluidity under hypoosmotic conditions but instead remained largely unchanged, paralleling the modest increases observed in run behavior (Figs. 5C-E).

**Fig. 5.**
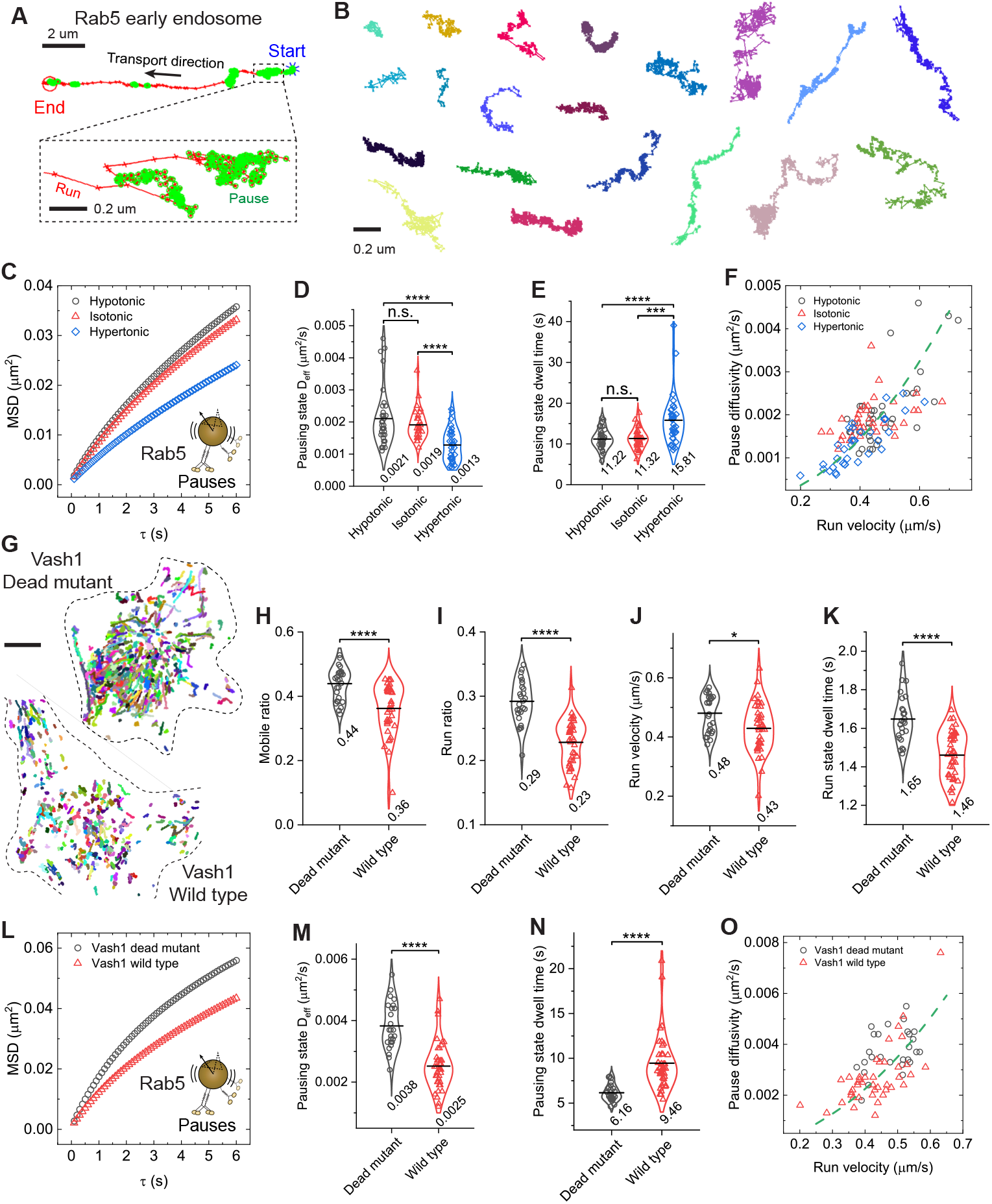
Motor engagement coordinates pause-run dynamics in dynein-driven transport of Rab5 early endosomes. (A) Magnified trajectory image showing a Rab5-positive early endosome from BEAS-2B cells. Blue stars and red circles mark the start and end point of the trajectory, respectively. Pause states are highlighted by green circles. Scale bar: 2 *µ*m. Inset shows a magnified view of the pause state. Scale bar: 200 nm. (B) Representative pausing segments extracted from mobile trajectories. Scale bar: 200 nm. (C) Measured MSDs of Rab5-positive early endosomes in the pausing state as a function of delay time *τ* for cells treated with different osmotic solutions. (D-E) Measured *D*_eff_ of Rab5-positive early endosomes in the pausing state (D) and pausing time (E) for cells treated with different osmotic solutions. \*\*\*\**p <*0.0001, \*\*\**p*=0.00011, n.s. indicates not significant. Total trajectories analyzed from the mobile fraction are n = 33130, 28305, and 25248 from n = 37, 34, and 36 cells from 3 independent experiments. (F) Measured pausing state diffusivity as a function of run-state velocity for Rab5-positive endosomes from each cell. The green dashed line represents a fit to a parabolic distribution curve, given by, *D*_eff_ = 0.009*v*^2^. (G) Representative trajectories of the mobile fraction of Rab5-positive endosomes for cells transiently expressing the wild type or the dead mutant of VASH1. Scale bars, 10 *µ*m. (H-K) Measured run parameters of Rab5-positive endosomes in BEAS-2B cells transiently expressing the wild type or the dead mutant of VASH1. \*\*\*\**p <*0.0001, \**p*=0.011. (L) Measured MSDs of Rab5-positive endosomes in the pausing state as a function of delay time *τ* for cells transiently expressing the wild type or the dead mutant of VASH1. (M-N) Measured *D*_eff_ of Rab5-positive endosomes in the pausing state (M) and pausing time (N) for cells transiently expressing the wild type or the dead mutant of VASH1. \*\*\*\**p <*0.0001. Total trajectories analyzed from the mobile fraction are n = 24849 and 27763 from n = 27 and 42 cells from 3 independent experiments. (O) Measured pausing state diffusivity as a function of run-state velocity for Rab5-positive endosomes from each cell. The green dashed line represents a fit to a parabolic distribution curve, given by, *D*_eff_ = 0.014*v*^2^. All violin graphs display all data points with means. P-values were calculated using a student’s t-test.

We then directly impaired motor-microtubule engagement. Since microtubule detyrosination inhibits the landing of dynein–dynactin complexes on microtubules via the p150 CAP-Gly domain in vitro (17, 50), we increased microtubule detyrosination by expressing VASH1. This efficiently impaired the run behavior of Rab5-positive endosomes (Figs. 5G–K). As a direct consequence, pausing-state diffusivity markedly decreased and trapping times during pauses increased (Figs. 5L–N). Across all perturbations, we observed a robust correlation between pause-state diffusivity and runstate velocity (Figs. 5F and 5O). Notably, this relationship for retrograde transport of Rab5-endosomes closely paralleled that observed for kinesin-driven transport of Rab6A-positive vesicles. When we combined all data from both cargo types, all points collapsed onto a single parabolic distribution curve (Fig. 6A). This demonstrates that the coordination of run and pause dynamics is a universal property of motor-driven transport, independent of motor type or direction, and is governed by the fundamental principle of motor engagement.

**Fig. 6.**
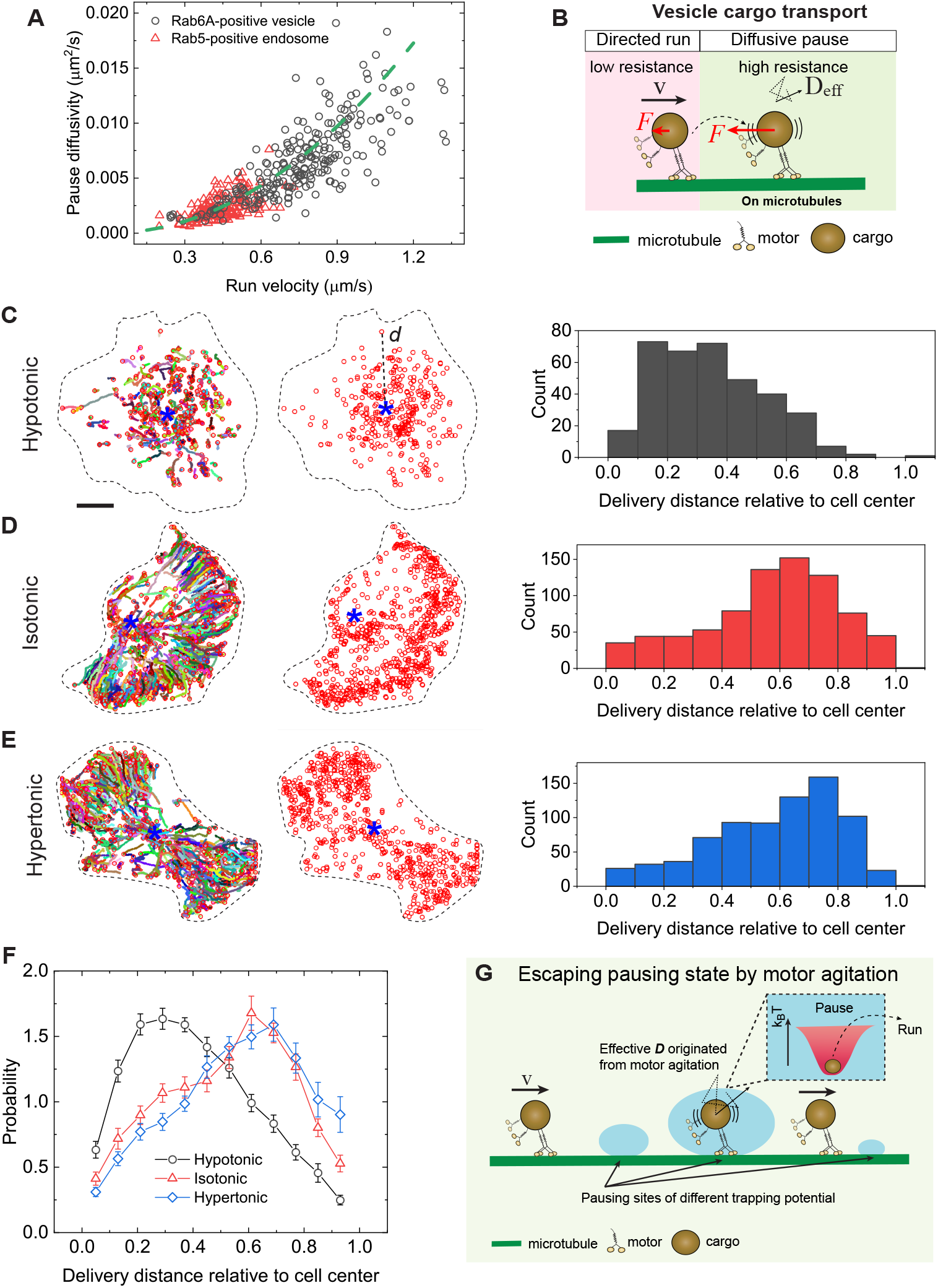
A molecular motor engaged pausing state facilitates the proper delivery of vesicle cargoes. (A) Measured pausing state diffusivity as a function of run-state velocity for Rab6A-positive vesicles and Rab5-positive endosomes in BEAS-2B cells across various perturbations. The green dashed line represents a fit to a parabolic distribution curve, given by, *D*_eff_ = 0.012*v*^2^. (B) Schematic illustration of the coordination between run and pause dynamics in intracellular transport mediated by motor proteins. Vesicle cargoes exhibit directed motility with a characteristic velocity (*v*) during the run state when the local resistance force (*F* ) is low, and diffusive-like motion with an apparent diffusivity (*D*_eff_) during the pause state when the local resistance is high. (C-E) Representative trajectories (colored lines, left) of the mobile fraction of Rab6A-positive vesicles for cells treated with different osmotic solutions. The end positions of each vesicle trajectory (red circles) and the relative cell centers (blue stars) are used to calculate the delivery distance *d* of each vesicle during the observation. The histograms of delivery distance for a representative cell in each condition are shown on the right. Scale bars, 10 *µ*m. (F) Measured PDFs of delivery distance for Rab6A-positive vesicles in BEAS-2B cells treated with different osmotic solutions. The distances are averaged from n = 28, 26 and 22 cells from 3 independent experiments. (G) Proposed motor-assisted escape model. During intracellular transport within a complex living cell, vesicles driven by multiple motor proteins encounter regions of increased resistance or “sticky zones” along their microtubule tracks, where they exhibit diffusive pausing behavior. The effective number of active motor proteins engaged with the vesicle and the microtubule determines both the apparent diffusion coefficient and the escape time from these traps. These sticky zones can be conceptualized as potential wells of varying depth, with motor activity supplying the kinetic energy necessary for vesicles to overcome these energy barriers and resume directed transport.

### Motor-mediated escape from pauses ensures efficient cargo delivery

The robust coupling between pause and run states raised a key question: are these truly distinct states, or two manifestations of a single process where motors navigate variable local resistance? When resistance is low, vesicles move freely, like cargo-free motors. When resistance is high, a tug-of-war between motor forces and the viscoelastic cytoplasm produces jiggling, diffusive-like motion around microtubules, trapping vesicles transiently as pauses (Fig. 6B). The number of engaged motors therefore governs a vesicle’s mobility and, even more importantly, its ability to escape these transient traps. Since vesicles spend the majority of their transit time in the pause state, an efficient escape rate is vital for delivery. To test this hypothesis, we analyzed the final delivery positions of Rab6A vesicles under osmotic conditions that differentially regulate motor activity (25). Rab6A-positive vesicles are typically transported from the perinuclear Golgi to the cell periphery (24, 25). Under hypoosmotic conditions, where motor activity and pause escape are inhibited, histograms of the distances between the cell center and vesicle final positions revealed that vesicles were concentrated near the midzone of the cell, approximately halfway to the periphery. In contrast, under iso- and hyperosmotic conditions, where motor engagement is higher, vesicles were successfully delivered to the periphery (Figs. 6C–F), demonstrating that effective escape from pause states is essential for proper cargo delivery.

## Discussion

Taken together, our work reveals a unifying quantitative principle governing intracellular transport: a parabolic relationship between pause diffusivity and run velocity (*D*_eff_ ∝ *v*^2^). This demonstrates that pausing and running are not independent motility states but dynamically coupled manifestations of motor engagement with microtubules in the complex cytoplasmic environment. This finding challenges the longstanding view that pauses represent cargo detachment from microtubules and passive diffusion (19, 20, 22, 28, 51, 52). Instead, our data strongly supports that pauses predominantly correspond to active, motor-engaged states where teams of motors continuously generate force against local resistance. This integrated run-pause dynamic enables vesicles to actively navigate a heterogeneous intracellular landscape, balancing directed motion with transient trapping.

The prolonged pause escape times under reduced motor activity (Figs. 2G, 3L, 4L and 5N), support model in which motors perform the work necessary for vesicles to escape local energy barriers (53, 54). We conceptualize these trapping sites as potential wells of varying depths and motor activity supplies the kinetic energy to overcome these barriers and resume directed runs (Fig. 6G).

Given that Rab6A-vesicles and Rab5-endosomes move predominantly unidirectionally (Figs. 1F and 5A) (25), and inhibiting kinesin-1 with kinesore did not increase pausing mobility or run ratio (Figs. 3E and 3K), the observed motor-engaged pausing is unlikely to represent the “draw” state characteristic of a tug-of-war between opposing motors (21, 28, 33, 55). This interpretation is reinforced by the nearly 3-fold difference in pausing mobility between Rab6Avesicles and Rab5-endosomes (Figs. 2F and 5D). If cargos were in a “draw” state and simply following fluctuating microtubules to which they are tethered, their pause mobilities would be similar. Instead, the resistance to motor-generated pulling forces may arise from transient tethering or obstruction by other intracellular structures (8, 22, 23, 32, 56). These structures could act as elastic elements, and the resulting force balance between motor pulling and this elastic resistance could produce the diffusive-like motion and observed parabolic scaling *D*_eff_ ∝ *v*^2^.

Cells also appear to actively adjust motor-generated forces to adapt to environmental changes. Under hyperosmotic stress, the 39% decrease in cytoplasmic fluidity (Fig. 2D) was matched by a 32% reduction in pausing mobility for Rab5endosomes, while Rab6A-vesicles showed no change (Fig. 2F). This discrepancy suggests that kinesin-driven transport may selectively upregulate force to counterbalance the elevated viscoelastic drag, potentially through enhanced recruitment of kinesin-1 by MAP7, which is enriched on microtubules under hypertonicity (10, 25, 56, 57). Conversely, under hypoosmotic stress, the 39% increase in fluidity (Fig. 2D) coincided with a striking 47% decrease in pausing mobility for Rab6A-vesicles (Fig. 2F), while Rab5-endosomes were unaffected (Fig. 5D). We previously showed that hypotonic treatment disassociates MAP7 from microtubules and increases microtubule detyrosination (25). The combined loss of the kinesin-1 recruitment by MAP7 and inhibition of kinesin-3/dynein via detyrosination likely underlies the strong suppression of motor activity and pausing mobility, overriding the changes in fluidity. This highlights how cells can tune transport by locally modulating the microtubule landscape, allowing motor-cargo complexes to adapt to a heterogeneously crowded cytoplasm.

## Conclusion

Overall, the robustness of the *D*_eff_ ∝ *v*^2^ scaling across cargo types, motor families, and perturbations suggests it is a fundamental physical property of motor teams navigating a resistive environment. We speculate this relationship could be a general feature of processive molecular motors operating in confined, viscoelastic cytoplasm. Future studies could test this model in vitro using optical traps to apply calibrated loads to beads driven by defined motor teams (57–59), measuring whether the parabolic relationship between velocity under low load and diffusivity under high load is reconstituted.

## MATERIALS AND METHODS

### Cell line

BEAS-2B cells (ATCC, CRL-9609) were maintained in Dulbecco’s modified Eagle’s medium (DMEM, Gibco), supplemented with 10% fetal bovine serum (FBS), 50 units/mL of penicillin and 50 *µ*g/mL of streptomycin. All cell cultures were maintained in a 95% air/5% CO_2_ atmosphere at 37 °C. The cell line was routinely confirmed to test negative for mycoplasma contamination. For live cell imaging, cells were seeded at a density of approximately 5×10^5^ cm^*−*2^ on a glass coverslip which was placed in a 35mm polystyrene tissue-culture dish. Transfections were performed by using the FuGENE 6 (Promega) according to manufacturer’s instructions. Cells were generally transfected for 8 hours with 1 µg plasmids when the density reached *∼* 80% confluency.

### Antibodies and immunostaining

For immunostaining, we used mouse monoclonal antibodies against *α*-tubulin (DM1A, T9026; Sigma) and rabbit polyclonal antibodies against detyrosinated *α*-tubulin (AB3201; Sigma). The following secondary antibodies were used: Alexa Fluor 488– and 647–conjugated goat antibodies against rabbit and mouse IgG (Invitrogen) for immunofluorescence. For immunofluorescence cell staining, BEAS-2B cells were generally fixed in *−* 20 °C methanol for 10 min, and then blocked with 4% bovine serum albumin (BSA, Sigma) in PBS at room temperature for 2 hours. Next, cells were incubated with primary antibodies against detyrosinated tubulin (1:200) and *α*-tubulin (1:500), and corresponding secondary antibodies (1:200), each for 1 hour. Stained cells were mounted onto a microscope slide with ProLong^*T M*^ Gold Antifade Mountant (Invitrogen), and then examined by using spinning disk confocal microscopy. The spinning disk confocal was performed on an inverted research microscope Eclipse Ti2-E with the Perfect Focus System (Nikon), equipped with a Plan Apo 60× NA 1.40 oil objective, a Crest X-Light V3 spinning disk confocal head (Crest-Optics), a Celesta light engine (Lumencor) as the light source, a Prime 95B 25MM sCMOS camera (Teledyne Photometrics) and controlled by NIS elements AR software (Nikon). The fluorescent images for BEAS-2B cells were collected over a stack of vertical z-sections across the entire cell *∼* 4 *µ*m thickness. The final fluorescent images and their fluorescent intensities shown in the main text are based on the z-averaged images by using Fiji software (https://fiji.sc/).

### Generation of plasmids

The expression vectors used in this study were pDeltaCMV-KIF5B(1-560)-mStayGold, pEGFP-Rab6A (Addgene plasmid 49469), pEGFP-Rab5 (Addgene plasmid 49888), pCMV-VASH1-EGFP-P2A-SVBP, pCMV-VASH1(C168A, catalytically dead mutant)-EGFP-P2A-SVBP, pCMV-VASH1-mScarlet-P2A-SVBP, pCMV-VASH1(C168A, catalytically dead mutant)-mScarlet-P2A-SVBP, pDeltaCMV-VASH1-mScarlet-P2A-SVBP, pDeltaCMV-VASH1(C168A, catalytically dead mutant)-mScarlet-P2A-SVBP and pCDNA3.1-pCMV-PfV-GS-Sapphire (Addgene plasmid 116933; for mammalian expression of 40 nm-GEMs). mStayGold was synthesized as a gBlock based on the published sequences (36). The truncated KIF5B proteins (a.a. 1-560) were cloned into a pDeltaCMV vector (a gift from Dr. Scott Hansen for expressing proteins at low levels (25, 37)) with a C-terminal mStayGold cassette. VASH1 and SVBP proteins were cloned into a pDeltaCMV vector or pCMV vector split by a P2A selfcleaving peptide with a EGFP or mScarlet cassette at the C-terminus of VASH1 (25, 26). The single amino acid mutations were generated by PCR. All cloning were performed using Gibson assembly. All constructs were verified by DNA sequencing (Plasmidsaurus).

### Osmotic and drug treatments

To manipulate the cytoplasm fluidity, cells were treated with and imaged in extracellular osmotic environments ranging from hypoosmotic to hyperosmotic conditions (250–400 mOsm). These solutions were prepared by adding, respectively, 0.28 (*∼* 400 mOsm, hypertonic), 0.2 (*∼* 310 mOsm, isotonic control), 0.15 (*∼* 250 mOsm, hypotonic) and 0.1 (*∼* 200 mOsm, hypotonic) D-mannitol to a hypotonic base solution (in millimolar: 40 NaCl, 5 KCl, 1 CaCl_2_, 2 MgCl_2_, 10 HEPES (pH 7.4), *∼* 91 mOsm) to maintain a constant ionic strength (25, 60). Cells were typically treated with the osmotic solutions 10 min before imaging. To activate kinesin-1 activity, cells were treated with kinesore (100 *µ*M, TORCIS) in culture medium for 1 hour before being fixed or for 10 min before being imaged using TIRF microscopy.

### Total internal reflection fluorescence microscopy

TIRF microscopy experiments were performed on an inverted research microscope Eclipse Ti2-E with the Perfect Focus System (Nikon), equipped with a 1.49 NA 100× TIRF objective with the 1.5× tube lens setting, a Ti-S-E motorized stage, piezo Z-control (Physik Instrumente), LU-N4 laser units (Nikon) as the light source, an iXon DU897 cooled EMCCD camera (Andor) with an high-speed emission filter wheel (ET480/40M for mTurquoise2, ET525/50M for GFP, ET520/40M for YFP, and ET632/60M for mRuby2; Chroma). The microscope was controlled with NIS Elements software (Nikon). All live-cell experiments were performed in a live-cell imaging chamber (H301-Nikon-TI-S-ER, Oko Labs) that was equipped to the microscope to provide optimal culture conditions (95% air/5% CO_2_ atmosphere at 37 °C) for cells during imaging. After cell transfection, the cell-containing glass coverslip was mounted on a coverslip holder (SC15012, Aireka Cells), which was finally mounted on the microscope. KIF5B(1-560) was recorded at 5 fps for 3 min. Rab6A and Rab5 vesicles were recorded at 10 fps for 3 min. 40-nm-GEMs were recorded at 20 fps for 2 min. The recorded images have 16 bits of gray scales and a spatial resolution of 512×512 pixels with the width of each pixel = 107 nm in our TIRF optical setup.

### Single particle tracking and analysis

Single particle tracking (SPT) was performed using a homemade tracking program written in Matlab as previously described (25, 61), which is based on the standard tracking algorithm (62, 63).

With this advanced SPT algorithm, we were able to obtain the position **r**(*t*) at time *t* for KIF5B motors, Rab6A-positive secretory vesicles, Rab5-positive early endosomes and GEMs, and their trajectories were constructed from the consecutive images. This algorithm allowed us to achieve a spatial tracking resolution of *∼* 20 nm.

To characterize the mixed-motions of Rab6A-positive secretory vesicles and Rab5-positive early endosomes, we used an algorithm that could automatically identify and extract the states of diffusive “jiggling” movement and the states of directed “runs”, as described previously (25, 64, 65). Briefly, we determined the motion state of an arbitrary point in the trajectory by analyzing the turning angles around it at 5 different time scales. Trajectories containing no directed runs were considered as immobile. The mobile ratio was defined as the ratio of the number of mobile trajectories over the total number of trajectories in a single cell. The run ratio was defined as the ratio of the total time spent in the run state over the total travel time of a vesicle. The mobile ratio and run ratio generally characterize how active the motors are in moving vesicles from the “pause” to the “run” states at the whole cell level. Directed runs longer than 20 time-steps (2 s) were used to compute the run velocity. This is achieved by computing the mean squared displacements (MSDs), *⟨Δ* **r**^2^(*τ* )*⟩* = *⟨*(**r**(*t* + *τ* ) *−* **r**(*t*))^2^*⟩*, and fitted to *⟨Δ* **r**^2^(*τ* )*⟩* = *v*^2^*τ* ^2^ to obtain the run velocity *v*. Diffusive pauses longer than 10 time-steps (1 s) were used to compute the effective pause diffusivity *D*_eff_. *D*_eff_ was obtained by fitting the MSDs to *⟨Δ* **r**^2^(*τ* )*⟩* = 4*D*_eff_*τ* . Run-state and pause-state dwell time of each trajectory were obtained by averaging the dwell time of each run and pause segments, respectively. To quantify the delivery distance for Rab6A positive vesicles, we calculated the distance *d* between the end position of each trajectory (*x, y*) to the cell center (*x*_0_, *y*_0_). Because microtubules in BEAS-2B cells are radially organized, we defined the cell center as the position where all vesicle transport started. In each cell, we normalized *d* as *d*_norm_ = (*d − d*_min_)*/*(*d*_max_ *− d*_min_), such that *d*_norm_ from cells of different sizes are in the same range 0 *≤ d*_norm_ *≤* 1. For the quantification of KIF5B motor run time, we only considered trajectories that lasted longer than 5 time-steps (1 s).

To study the diffusion dynamics of GEMs, we first selected the mobile trajectories from the whole set of GEMs trajectories. This is achieved by computing the radius of gyration *R*_*g*_(*τ* ) of each GEMs trajectory obtained over a time period of *τ*,

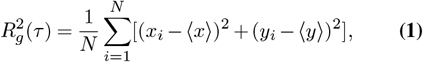

where *N* is the total number of time steps in each trajectory, *x*_*i*_ and *y*_*i*_ are the projection of the position of each trajectory step on the *x−* and *y−* axis, respectively, and *⟨x⟩* and *⟨y ⟩* are their mean values. Physically, *R*_*g*_ quantifies the size of a GEMs trajectory generated during the time lapse *τ* . A cutoff value of 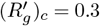 was used in the experiment, below which the GEMs trajectories are treated as immobile ones (25, 61). Here 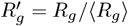 is the normalized radius of gyration, with *R*_*g*_ being the mean value of *⟨ R*_*g*_*⟩*. MSDs of individual mobile trajectories were then computed and fitted to *⟨Δ* **r**^2^(*τ* )*⟩* = 4*D*_eff_*τ* to obtain the trajectory-based effective diffusion coefficients of GEMs, *D*_eff_, in BEAS-2B cells.

### Statistics

Data are expressed as mean ± s.e.m. unless specified otherwise. Graphs were created using Origin. Statistical tests were performed with two-tailed unpaired Student’s t-test. The statistical details of each experiment can be found in the figure legends.

## Acknowledgments

The authors wish to thank all the members of the Ori-McKenney and McKenney laboratories for their kind help and feedback. This work is supported by the NIH grant 1R35GM133688 to K.M.O.-M.

## Author Contributions

Y. S. and K. M. O. M. conceived the project and designed the experiments. Y. S. performed all experiments and analyzed the data. Y. S. and K. M. O. M. wrote the manuscript.

## Declaration of Interests

The authors declare no competing financial interests.

## Lead contact

Further information and requests for resources and reagents should be directed to and will be fulfilled by the lead contact, Kassandra M. Ori-McKenney.

## Materials availability

Cell lines and plasmids are available upon request from the authors.

## Data and code availability

Data generated and Matlab code used in this study are available upon request.

## Supplementary Figures

**Fig. S1.**
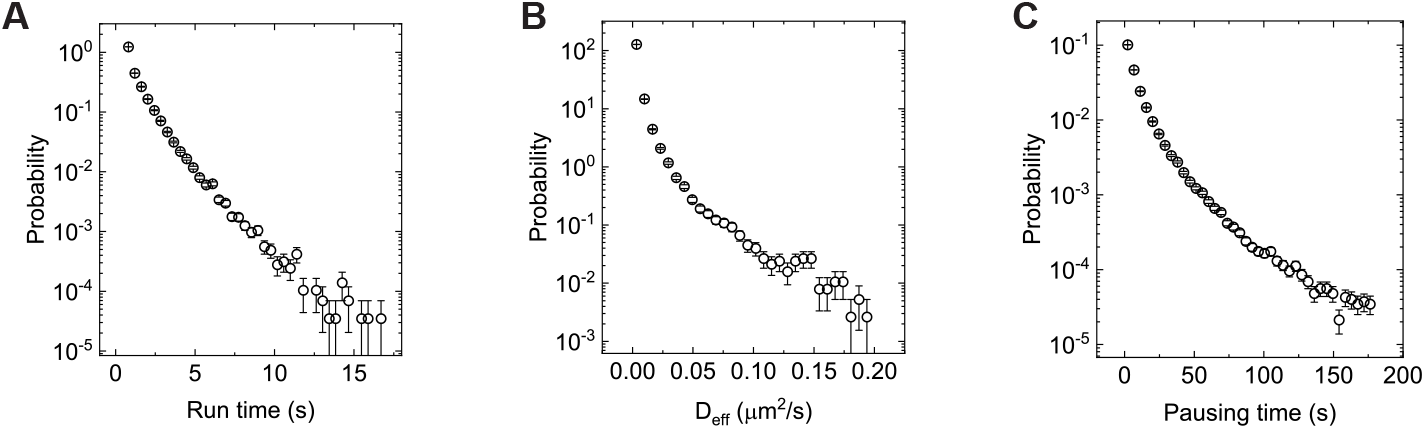
Pausing state of Rab5-positive endosome transport is vastly heterogeneous. (A) Measured PDF of the dwell time for run segments extracted from mobile trajectories. Total segments analyzed are n = 70451. (B and C) Measured PDF of the effective diffusion coefficient *D*_eff_ (B) and dwell time (C) in the pausing state for Rab5-positive endosomes. Total pausing segments analyzed from the mobile fraction are n = 84165.

## Notes

### Competing Interest Statement

The authors have declared no competing interest.

## REFERENCES

1. Vale, R. D. The molecular motor toolbox for intracellular transport. Cell 112, 467–480 (2003).

2. Mogre, S. S., Brown, A. I. & Koslover, E. F. Getting around the cell: physical transport in the intracellular world. Phys. Biol. 17, 061003 (2020).

3. Brangwynne, C. P., Koenderink, G. H., MacKintosh, F. C. & Weitz, D. A. Intracellular transport by active diffusion. Trends Cell Biol. 19, 423–427 (2009).

4. Bonucci, M., Shu, T. & Holt, L. J. How it feels in a cell. Trends Cell Biol. (2023).

5. Moeendarbary, E. et al. The cytoplasm of living cells behaves as a poroelastic material. Nat. Mater. 12, 253–261 (2013).

6. Ross, J. L., Shuman, H., Holzbaur, E. L. & Goldman, Y. E. Kinesin and dynein-dynactin at intersecting microtubules: motor density affects dynein function. Biophys. J. 94, 3115–3125 (2008).

7. Voeltz, G. K., Sawyer, E. M., Hajnóczky, G. & Prinz, W. A. Making the connection: How membrane contact sites have changed our view of organelle biology. Cell 187, 257–270 (2024).

8. Jongsma, M. L. et al. An ER-associated pathway defines endosomal architecture for controlled cargo transport. Cell 166, 152–166 (2016).

9. Bodakuntla, S., Jijumon, A. S., Villablanca, C., Gonzalez-Billault, C. & Janke, C. Microtubule-associated proteins: structuring the cytoskeleton. Trends Cell Biol. 29, 804–819 (2019).

10. Monroy, B. Y., Sawyer, D. L., Ackermann, B. E., Borden, M. M., Tan, T. C. & Ori-McKenney, K. M. Competition between microtubule-associated proteins directs motor transport. Nat. Commun. 9, 1487 (2018).

11. Monroy, B. Y. et al. A combinatorial MAP code dictates polarized microtubule transport. Dev. Cell 53, 60–72 (2020).

12. Vershinin, M., Carter, B. C., Razafsky, D. S., King, S. J. & Gross, S. P. ultiple-motor based transport and its regulation by Tau. Proc. Natl Acad. Sci. 104, 87–92 (2007).

13. Tan, R. et al. Microtubules gate tau condensation to spatially regulate microtubule functions. Nat. Cell Biol. 21, 1078–1085 (2019).

14. Hammond, J. W., Cai, D. & Verhey, K. J. Tubulin modifications and their cellular functions. Curr. Opin. Cell Biol. 20, 71–76 (2008).

15. Janke, C. & Magiera, M. M. The tubulin code and its role in controlling microtubule properties and functions. Nat. Rev. Mol. Cell Biol. 21, 307–326 (2020).

16. Sirajuddin, M., Rice, L. M. & Vale, R. D. Regulation of microtubule motors by tubulin isotypes and post-translational modifications. Nat. Cell Biol. 16, 335–344 (2014).

17. McKenney, R. J., Huynh, W., Vale, R. D. & Sirajuddin, M. Tyrosination of α-tubulin controls the initiation of processive dynein–dynactin motility. EMBO J. 35, 1175–1185 (2016).

18. Fan, X. & McKenney, R. J. Control of motor landing and processivity by the CAP-Gly domain in the KIF13B tail. Nat. Commun. 14, 4715 (2023).

19. Gross, S. P., Welte, M. A., Block, S. M. & Wieschaus, E. F. Dynein-mediated cargo transport in vivo: a switch controls travel distance. J. Cell Biol. 148, 945–956 (2000).

20. Welte, M. A. Bidirectional transport along microtubules. Curr. Biol. 14, R525–R537 (2004).

21. Hendricks, A. G., Perlson, E., Ross, J. L., Schroeder, H. W., Tokito, M. & Holzbaur, E. L. Motor coordination via a tug-of-war mechanism drives bidirectional vesicle transport. Curr. Biol. 20, 697–702 (2010).

22. Zajac, A. L., Goldman, Y. E., Holzbaur, E. L. & Ostap, E. M. Local cytoskeletal and organelle interactions impact molecular-motor-driven early endosomal trafficking. Curr. Biol. 23, 1173–1180 (2013).

23. Bálint, Š., Verdeny Vilanova, I., Sandoval Álvarez, Á. & Lakadamyali, M. Correlative live-cell and superresolution microscopy reveals cargo transport dynamics at microtubule intersections. Proc. Natl Acad. Sci. 110, 3375–3380 (2013).

24. Serra-Marques, A. et al. Concerted action of kinesins KIF5B and KIF13B promotes efficient secretory vesicle transport to microtubule plus ends. Elife 9, e61302 (2020).

25. Shen, Y. & Ori-McKenney, K. M. Microtubule-associated protein MAP7 promotes tubulin posttranslational modifications and cargo transport to enable osmotic adaptation. Dev. Cell 59, 1553–1570 (2024).

26. Shen, Y., Maxson, R., McKenney, R. J. & Ori-McKenney, K. M. Microtubule acetylation is a biomarker of cytoplasmic health during cellular senescence. bioRxiv 2025–03 (2025).

27. Sheng, Z. H. Mitochondrial trafficking and anchoring in neurons: new insight and implications. J. Cell Biol. 204, 1087–1098 (2014).

28. Hancock, W. O. Bidirectional cargo transport: moving beyond tug of war. Nat. Rev. Mol. Cell Biol. 15, 615–628 (2014).

29. McKenney, R. J., Huynh, W., Tanenbaum, M. E., Bhabha, G. & Vale, R. D. Activation of cytoplasmic dynein motility by dynactin-cargo adapter complexes. Science 345, 337–341 (2014).

30. Chiba, K., Ori-McKenney, K. M., Niwa, S. & McKenney, R. J. Synergistic autoinhibition and activation mechanisms control kinesin-1 motor activity. Cell Rep. 39, 110900 (2022).

31. van Bommel, B., Konietzny, A., Kobler, O., Bär, J. & Mikhaylova, M. F-actin patches associated with glutamatergic synapses control positioning of dendritic lysosomes. EMBO J. 38, e101183 (2019).

32. Kang, J. S., Tian, J. H., Pan, P. Y., Zald, P., Li, C., Deng, C. & Sheng, Z. H. Docking of axonal mitochondria by syntaphilin controls their mobility and affects short-term facilitation. Cell 132, 137–148 (2008).

33. D’Souza, A. I., Grover, R., Monzon, G. A., Santen, L. & Diez, S. Vesicles driven by dynein and kinesin exhibit directional reversals without regulators. Nat. Commun. 14, 7532 (2023).

34. Mohan, N., Sorokina, E. M., Verdeny, I. V., Alvarez, A. S. & Lakadamyali, M. Detyrosinated microtubules spatially constrain lysosomes facilitating lysosome–autophagosome fusion. J. Cell Biol. 218, 632–643 (2019).

35. Taylor, E. W. & Borisy, G. G. Kinesin processivity. J. Cell Biol. 151, F27–F30 (2000).

36. Ivorra-Molla, E. et al. A monomeric StayGold fluorescent protein. Nat. Biotechnol. 42, 1368–1371 (2024).

37. Watanabe, N. & Mitchison, T. J. Single-molecule speckle analysis of actin filament turnover in lamellipodia. Science 295, 1083–1086 (2002).

38. Jacobson, C., Schnapp, B. & Banker, G. A. A change in the selective translocation of the Kinesin-1 motor domain marks the initial specification of the axon. Neuron 49, 797–804 (2006).

39. Cai, D., Verhey, K. J. & Meyhöfer, E. Tracking single Kinesin molecules in the cytoplasm of mammalian cells. Biophys. J. 92, 4137–4144 (2007).

40. Cai, D., McEwen, D. P., Martens, J. R., Meyhofer, E. & Verhey, K. J. Single molecule imaging reveals differences in microtubule track selection between Kinesin motors. PLoS Biol. 7, e1000216 (2009).

41. Schlager, M. A. et al. Bicaudal d family adaptor proteins control the velocity of Dynein-based movements. Cell Rep. 8, 1248–1256 (2014).

42. Guo, M. et al. Probing the stochastic, motor-driven properties of the cytoplasm using force spectrum microscopy. Cell 158, 822–832 (2014).

43. Molines, A. T. et al. Physical properties of the cytoplasm modulate the rates of microtubule polymerization and depolymerization. Dev. Cell 57, 466–479 (2022).

44. Randall, T. S. et al. A small-molecule activator of kinesin-1 drives remodeling of the microtubule network. Proc. Natl Acad. Sci. 114, 13738–13743 (2017).

45. Andreu-Carbó, M., Fernandes, S., Velluz, M. C., Kruse, K. & Aumeier, C. Motor usage imprints microtubule stability along the shaft. Dev. Cell 57, 5–18 (2022).

46. Brangwynne, C. P., Koenderink, G. H., MacKintosh, F. C. & Weitz, D. A. Cytoplasmic diffusion: molecular motors mix it up. J. Cell Biol. 183, 583–587 (2008).

47. Aillaud, C. et al. Vasohibins/SVBP are tubulin carboxypeptidases (TCPs) that regulate neuron differentiation. Science 358, 1448–1453 (2017).

48. Ramirez-Rios, S. et al. VASH1–SVBP and VASH2–SVBP generate different detyrosination profiles on microtubules. J. Cell Biol. 222, e202205096 (2022).

49. Flores-Rodriguez, N., Rogers, S. S., Kenwright, D. A., Waigh, T. A., Woodman, P. G. & Allan, V. J. Roles of dynein and dynactin in early endosome dynamics revealed using automated tracking and global analysis. PLOS ONE 6, e24479 (2011).

50. Nirschl, J. J., Magiera, M. M., Lazarus, J. E., Janke, C. & Holzbaur, E. L. α-Tubulin tyrosination and CLIP-170 phosphorylation regulate the initiation of dynein-driven transport in neurons. Cell Rep. 14, 2637–2652 (2016).

51. Kaplan, L., Ierokomos, A., Chowdary, P., Bryant, Z. & Cui, B. Rotation of endosomes demonstrates coordination of molecular motors during axonal transport. Sci. Adv. 4, e1602170 (2018).

52. Gennerich, A., Carter, A. P., Reck-Peterson, S. L. & Vale, R. D. Force-induced bidirectional stepping of cytoplasmic dynein. Cell 131, 952–965 (2007).

53. Hänggi, P., Talkner, P. & Borkovec, M. Reaction-rate theory: fifty years after Kramers. Rev. Mod. Phys. 62, 251 (1990).

54. Sharma, A., Wittmann, R. & Brader, J. M. Escape rate of active particles in the effective equilibrium approach. Phys. Rev. E 95, 012115 (2017).

55. Kunwar, A. et al. Mechanical stochastic tug-of-war models cannot explain bidirectional lipiddroplet transport. Proc. Natl Acad. Sci. 108, 18960–18965 (2011).

56. Kunwar, A., Vershinin, M., Xu, J. & Gross, S. P. Stepping, strain gating, and an unexpected force-velocity curve for multiple-motor-based transport. Curr. Biol. 18, 1173–1183 (2008).

57. Kunwar, A. & Mogilner, A. Robust transport by multiple motors with nonlinear force–velocity relations and stochastic load sharing. Phys. Biol. 7, 016012 (2010).

58. Jamison, D. K., Driver, J. W. & Diehl, M. R. Cooperative responses of multiple kinesins to variable and constant loads. J. Biol. Chem. 287, 3357–3365 (2012).

59. Rao, L. et al. The Power of Three: Dynactin associates with three dyneins under load for greater force production. bioRxiv 2025-01 (2025).

60. Shen, Y. et al. Mechanical characterization of microengineered epithelial cysts by using atomic force microscopy. Biophys. J. 112, 398–409 (2017).

61. Shen, Y. et al. Directed motion of membrane proteins under an entropy-driven potential field generated by anchored proteins. Phys. Rev. Res. 3, 043195 (2021).

62. Crocker, J. C. & Grier, D. G. Dethods of digital video microscopy for colloidal studies. J. Colloid Interface Sci. 179, 298–310 (1996).

63. Anthony, S., Zhang, L. & Granick, S. Methods to track single-molecule trajectories. Langmuir 22, 5266–5272 (2006).

64. Shen, Y., Wen, Y., Zhao, Q., Huang, P., Lai, P.Y. & Tong, P. Endosome-ER Interactions Define a Cellular Energy Landscape to Guide Cargo Transport. bioRxiv 2023-06 (2023).

65. Röding, M., Guo, M., Weitz, D. A., Rudemo, M. & Särkkä, A. Identifying directional persistence in intracellular particle motion using Hidden Markov Models. Math. Biosci. 248, 140–145 (2014).

